# Characterization of early Alzheimer’s-like pathological alterations in non-human primates with aging: a pilot study

**DOI:** 10.1101/2021.07.21.453246

**Authors:** Hannah Jester, Saahj Gosrani, Huiping Ding, Xueyan Zhou, Mei-Chuan Ko, Tao Ma

## Abstract

Sporadic or late onset Alzheimer’s disease (LOAD) is a multifactorial neurodegenerative disease with aging the most known risk factor. Non-human primates (NHPs) may serve as an excellent model to study LOAD because of their close similarity to humans in many aspects including neuroanatomy and neurodevelopment. Recent studies reveal AD-like pathology in old NHPs. In this pilot study, we took advantage of brain samples from 6 Cynomolgus macaques that were divided into two groups: middle aged (average age 14.81 years) and older (average age 19.33 years). We found multiple AD-like pathological alteration in the prefrontal cortex (but not in the hippocampus) of the older NHPs including tau hyperphosphorylation, increased activity of AMP-activated protein kinase (AMPK), decreased expression of protein phosphatase 2A (PP2A), impairments in mitochondrial morphology and postsynaptic densities (PSDs) formation. These findings may provide insights into the factors contributing to the development of LOAD, particularly during the early stage transitioning from middle to old age. Future endeavors are warranted to elucidate mechanisms underlying the regional (and perhaps cellular) vulnerability with aging and the functional correlation of such pathological changes in NHPs.

## 1. Introduction

Alzheimer’s disease (AD) is the most common cause of dementia globally and a devastating neurodegenerative disease without effective disease modifying therapies available (Association, 2019; Barker et al., 2002; Lane et al., 2018). Vast majority of AD are sporadic or late onset AD (LOAD), compared to early onset familial AD (FAD) that accounts for less than 10% of all AD cases (Cacace et al., 2016). Canonical AD neuropathology includes buildup of extracellular amyloid (Aβ) plaques and intracellular tau neurofibrillary tangles (NFTs) (Deture and Dickson, 2019; Lane et al., 2018). Meanwhile, mounting evidence indicates a host of other pathophysiological changes that may play crucial roles in AD pathogenesis including dysregulation of energy metabolism and mitochondrial function, inflammation, oxidative stress, and synaptic dysfunction (Baglietto-Vargas et al., 2016; Biessels and Reagan, 2015; de la Monte and Tong, 2014; Ma and Klann, 2012; Tramutola et al., 2017; Wang, Xin et al., 2019). It is recognized that before dementia and cognitive impairment can be clinically diagnosed, AD is characterized by a long asymptomatic preclinical phase, during which many of the aforementioned pathological alterations have occurred (C. Vickers et al., 2016; De Strooper and Karran, 2016). Consistently, much efforts have been recently dedicated to define and characterize the preclinical or early (earliest) stages of AD, which would help understand the disease etiology and consequently identification of novel therapeutic targets and biomarkers for diagnosis/prognosis of the disease (C. Vickers et al., 2016; Mufson et al., 2016; Wang, X. et al., 2020).

Synapse loss is strongly correlated with the AD-associated cognitive decline (DeKosky et al., 1996; Koffie et al., 2011). Synaptic damage can be caused by a number of factors such as Aβ oligomers, tau hyperphosphorylation, and mitochondrial dysfunction (Guerrero-Muñoz et al., 2015; Ma et al., 2011; Ma and Klann, 2012; Su et al., 2008). It is established that mitochondrial dysfunction occurs with aging and is one of the early features of AD pathophysiology. With aging and in AD, mitochondria exhibits decreased electron transport chain (ETC) activity and decreased expression of mitochondrial proteins (Boffoli et al., 1994; Navarro and Boveris, 2007).

AMP-activated protein kinase (AMPK) is a cellular energy sensory that plays a critical role in maintaining energy homeostasis by regulating anabolic and catabolic cellular processes (Hardie, 2011; Hardie et al., 2012). Activity of AMPK is increased by phosphorylation of its catalytic α subunit (AMPKα) at Thr172, which involves upstream kinases including liver kinase B 1 (LKB1) and Calcium/Calmodulin dependent protein kinase kinase 2 (CAMKK2)(Yan et al., 2018). De-phosphorylation of AMPKα, which is mainly regulated by protein phosphatase 2A (PP2A), reduces AMPK activity (Salminen et al., 2016). Accumulating evidence indicates that abnormally increased AMPK activity contributes to many aspects of AD pathophysiology including Aβ production, tau phosphorylation, synaptic failure and cognitive deficits (Domise et al., 2016; Domise et al., 2019; Ma et al., 2014; Vingtdeux et al., 2011; Vingtdeux et al., 2010). In addition, AMPK can regulate mitochondrial biogenesis, dysregulation of which is characterized in the early stage of many neurodegenerative diseases including AD (Beal, 2007; Lin and Beal, 2006; Wang, W. et al., 2020).

Non-human primates (NHPs) are excellent models to study molecular mechanisms and therapeutic targets associated with human neuronal diseases because of their close similarity to humans in many aspects including neuroanatomy, physiology, development and cognitive function (Colman, 2018; Li et al., 2019). Cynomolgus macaques and other old world monkeys are our closest phylogenetic relatives and share similar structural and physiological changes in the aging process. NHPs also share sequence homology with multiple key AD proteins such as APP and tau (Arnsten et al., 2019). Old monkeys naturally develop AD-like pathology such as diffuse and dense-cored Aβ plaques, as well as hyperphosphorylated and paired helical filament (PHF) tau, which contribute to the NFT development (Latimer et al., 2019; Nakamura et al., 1998; Oikawa et al., 2010; Wu et al., 2008). Thus, NHPs hold a great potential for deciphering early pathophysiological changes in aging and AD-like conditions. Here, we carried out pilot studies to characterize AD-like pathology and potential molecular signaling dysregulation in a non-human primate model during the transition from middle age to old age, which has rarely been investigated before to our knowledge.

## 2. Materials and Methods

### 2.1. Animals

All non-human primates were individually housed at the Wake Forest University Primate Center, on a 12hr light/dark cycle, and given regular chow (5038; LabDiet, St. Louis, MO, USA). They were provided fruit and water ad *libitum*. All experimental procedures were performed in accordance with the Guide for the Care and Use of Laboratory Animals and approved by the Institutional Animal Care and Use Committee of Wake Forest University School of Medicine (Winston-Salem, NC, USA). Three Cynomolgus macaques (*Macaca fascicularis*), 2 females and 1 male, were used as middle-aged controls. Three Cynomolgus macaques, all female, were used in the older group with significant age difference. Demographic Information is listed in Table 1.

**Table 1:** Cynomolgus Demographic Data

| Table 1: Cynomolgus Demographic Data |  |  |  |  |  |  |  |  |
| --- | --- | --- | --- | --- | --- | --- | --- | --- |
|  | Clinical ID | Gender | Age (yrs) | Body Weight (kg) | Brain Weight (g) | Glucose (mg/dL) | Cholesterol (mg/dL) | Triglycerides (mg/dL) |
| Middle-Aged | 7405/37 | Male | 15.08 | 11.48 | 74.65 | 55 | 88 | 131 |
|  | 7406/37 | Female | 14.67 | 6.31 | 64.97 | 75 | 137 | 62 |
|  | 7430/37 | Female | 14.67 | 7.18 | 68.00 | 69 | 117 | 88 |
|  |  | Mean | 14.81 | 8.32 | 69.21 | 66.33 | 114.00 | 93.67 |
|  |  | SEM | 0.14 | 1.60 | 2.86 | 5.93 | 14.22 | 20.12 |
| Older | 7871/35 | Female | 19.17 | 3.30 | 58.37 | 84 | 153 | 159 |
|  | 7676/37 | Female | 19.92 | 4.00 | 66.30 | 87 | 114 | 102 |
|  | 7675/37 | Female | 18.91 | 6.30 | 55.98 | 126 | 96 | 158 |
|  |  | Mean | 19.33*** | 4.53 | 60.22 | 99.00 | 121.00 | 139.67 |
|  |  | SEM | 0.30 | 0.91 | 3.12 | 13.53 | 16.82 | 18.84 |

### 2.2. Fixed Tissue Preparation

Tissue was dissected from appropriate locations and fixed in 4% paraformaldehyde for three days and transferred to a 30% sucrose cryoprotectant solution for 1 week. The tissue was then flash-frozen in a dry ice bath of 2-methylbutane and stored at −80C.

### 2.3. Immunohistochemistry

The tissue samples were equilibrated to the cryostat (HM525NX, ThermoFisher) temperature (−20C) for at least an hour. The tissue was then transferred to a pre-labeled plastic cryomold and entirely embedded with OCT (optimizing cutting temperature) and allowed to freeze completely inside the cryostat. The OCT embedded tissue block was then removed from the cryomold and immediately placed on a room temperature specimen chuck with a thin layer of OCT. The chuck was chilled inside the cryostat on the freeze rail, and after complete solidification, the specimen chuck with the tissue block adhered was attached to the specimen holder in the cryostat. Tissue was sectioned at a desired thickness (5-10um) and placed on SuperFrost Plus Slides. If staining with Hematoxylin (7211, Richard Allan) and Eosin (7111, Richard Allan), slides were immersed in 10% neutral buffered formalin as a first step and then stained on an automatic stainer (Gemini AS Automated Slide Stainer, ThermoFisher).

Slides were allowed to come to room temperature before antigen retrieval was performed using citrate buffer (pH 6.0), heated to boiling in a microwave and boiled for 15 minutes. Endogenous peroxidase activity was blocked using 3% H2O2 for 25 min. Slides were incubated in a humidified chamber in primary antibody for Amyloid-Beta (6E10) (1:1000; Sigma, Cat#Sig39320) or p-Tau (AT8) (1:1000; Thermofisher, Cat#MN1020), overnight at 4°C. Sections were then incubated in biotinylated α mouse (6E10) or rabbit (AT8) secondary antibody (1:200; Vector Labs, Burlingame, CA) for 2 hrs at room temperature followed by Vectastain Elite ABC reagent (Vector Labs) for another 30 minutes. Primary and secondary antibodies as well as ABC reagent were diluted in 1% BSA/PBS. Vectastain Elite ABC kit Diaminobenzidine (DAB) reagent (Vector Labs) was diluted in DAB diluent (Vector Labs) for a working DAB solution. Sections were developed in DAB for 1 min. Slides were counterstained using Mayer’s hematoxylin for 8 mins. In between each step of immunohistochemistry, sections were rinsed using distilled water or PBS (pH 7.4). Negative controls were incubated in 1% BSA with no primary antibody. Sections were dehydrated in an alcohol series and cleared with xylene, cover-slipped, and dried overnight. Imaging was performed using BZX710 all-in-one fluorescent microscope (Keyence, Japan).

### 2.4. Western Blotting

Tissues were removed from appropriate structures and flash frozen on dry ice. Tissues were then homogenized in an appropriate lysis buffer and quantified as previously described (Zimmermann et al., 2019). Samples were loaded on 4-15% TGX™ Precast Gels (Bio-Rad), and transferred to nitrocellulose membranes. Membranes were blocked and then probed overnight at 4°C using primary antibodies of interest. Blots were washed and HRP-labeled secondary antibodies were added. Primary antibodies used: p-AMPKα (1:1000; Thr172) (Cell Signaling, Cat#2535), AMPKα (1:1000; Cell Signaling, Cat#5832), p-Tau (1:1000; AT8;Thermofisher, Cat#MN1020), p-Tau (1:500; Thr231; ThermoFisher, Cat#MN1040), p-Tau (1:1000; Ser396; Invitrogen, Cat#44-752G), Tau (1:1000; Invitrogen, Cat#AHB0042), p-GSK3β (1:1000; S9) (Cell Signaling, Cat#5558), GSK3β (1:1000; Cell Signaling, Cat#12456), PP2AC (1:1000; Cell Signaling, Cat#2038), and GAPDH (1:10,000, Cell Signaling, Cat#5174), p-LKB1 (1:1000; Ser428) (Cell Signaling, Cat#), LKB1 (1:1000; Cell Signaling, Cat#), p-ACC1 (1:1000; Ser79) (Cell Signaling, Cat#), ACC1 (1:1000; Cell Signaling, Cat#), p-ULK1 (1:1000; Ser317) (Cell Signaling, Cat#), ULK1 (1:1000; Cell Signaling, Cat#), PGC-1α (1:1000; Cell Signaling, Cat#), Mitofusin2 (1:1000; Cell Signaling, Cat#), DRP1 (1:1000; Cell Signaling, Cat#), OXPHOS Cocktail (1:1000; Cell Signaling, Cat#), SOD1 (1:1000; Cell Signaling, Cat#), SOD2 (1:1000; Cell Signaling, Cat#), Catalase (1:1000; Cell Signaling, Cat#), p-mTOR (1:1000; Ser2448) (Cell Signaling, Cat#), mTOR (1:1000; Cell Signaling, Cat#), p-S6K1 (1:1000; Ser389) (Cell Signaling, Cat#), S6K1 (1:1000; Cell Signaling, Cat#), p-4E-BP1 (1:1000; Thr37, Thr46) (Cell Signaling, Cat#), 4E-BP1 (1:1000; Cell Signaling, Cat#), p-eEF2 (1:1000; Thr56) (Cell Signaling, Cat#), eEF2 (1:1000; Cell Signaling, Cat#), PSD-95 (1:1000; Cell Signaling, Cat#), Synapsin2 (1:1000; Cell Signaling, Cat#),. All antibodies were diluted in either 5% wt/vol Milk/TBST or 5% wt/vol BSA/TBST. The blots were visualized using chemiluminescence (Clarity™ ECL; Bio-Rad) and the Bio-Rad ChemiDoc™ MP Imaging System. Densitometric analysis was performed using ImageJ software. Data were normalized to GAPDH (for total protein analysis) or relevant total proteins (for phospho-protein analysis) unless otherwise specified.

### 2.5. Transmission Electron Microscopy (TEM)

The CA1 of the hippocampus of previously fixed tissue (described above) was microdissected and were washed in buffer and post-fixed with 1% osmium tetroxide in phosphate buffer for an hour. Samples were then dehydrated through a graded series of ethanol solutions. For preparation of resin infiltration, the samples were incubated in propylene oxide for two 15-minute changes. The samples were subsequently infiltrated with 1:1, 1:2, and pure solutions of Spurr’s resin and placed in a 70°C oven overnight to cure. A Reichert-Jung Ultracut E ultramicrotome was used to obtain 90 nm thick sections, stained with lead citrate and uranyl acetate, and viewed with a Tecnai Spirit transmission electron microscope operating at 80 kV (FEI Co.). Images were obtained with a 2Vu CCD camera (Advanced Microscopy Techniques) at ×11,000 magnification. Analysis was performed using ImageJ as previously described (Ostroff et al., 2002; Ostroff et al., 2018). Imaging and analysis were done by investigators blinded to animal groups.

For the mitochondria analysis: Image J was used to determine the area and length (longest points across mitochondrion) (Martins et al., 2017). The inner/outer mitochondrial membranes were given a binary rating of if they were intact and the cristae morphology was given a binary intactness rating based on literature (Arrázola and Inestrosa, 2015; Arrázola et al., 2015). Imaging and analysis were done by investigators blinded to animal groups.

### 2.6. Statistical Analysis

Data are presented as mean +/- SEM. For comparisons between groups, a two-tailed unpaired student’s t-test was used. Error probabilities of *p*<0.05 were considered statistically significant. Outliers were determined via Grubbs test. Statistics were performed using Prism 8 statistics software (GraphPad Software, San Diego, CA).

## 3. Results

### 3.1. Age-related increase of tau phosphorylation in the prefrontal cortex of Cynomolgus macaques

We first examined potential alterations on Aβ and tau pathology in different brain regions by performing immunohistochemical experiments. In the prefrontal cortex (PFC) of older NHPs, we observed a trending (p=0.0996) increase of Aβ (using the 6E10 antibody) staining and significant increase of tau phosphorylation (using the AT8 antibody), compared to the middle-aged group (Fig. 1A). We did not find any age-related alteration of either Aβ or tau pathology in the hippocampus (Fig. 1B). Interestingly, most Aβ staining (for either PFC or hippocampus) appears to be localized intra-neuronally. Using a biochemical approach, we further investigated potential age-related change of tau phosphorylation. Previous studies indicate that phosphorylation of tau at Thr231 has been found largely in PHF and pre-tangle formations, while Ser396 phosphorylation has been mostly associated with late-stage NFTs (Augustinack et al., 2002; Kimura et al., 1996; Neddens et al., 2018). We did not find age-related change in tau phosphorylation at the Ser396 site in either PFC or hippocampus (Fig. 1C and D). In comparison, levels of tau phosphorylation at the Thr231 site were significantly increased in the PFC of the older monkeys compared to the middle-age group (Fig. 1C). The hippocampi of the older group did not display any age-related changes in tau phosphorylation at the Thr231 site, compared to the middle-aged group (Fig. 1D). Furthermore, tau phosphorylation at the Thr231 site in the PFC is correlated with age (Fig. 1E).

**Figure 1.**
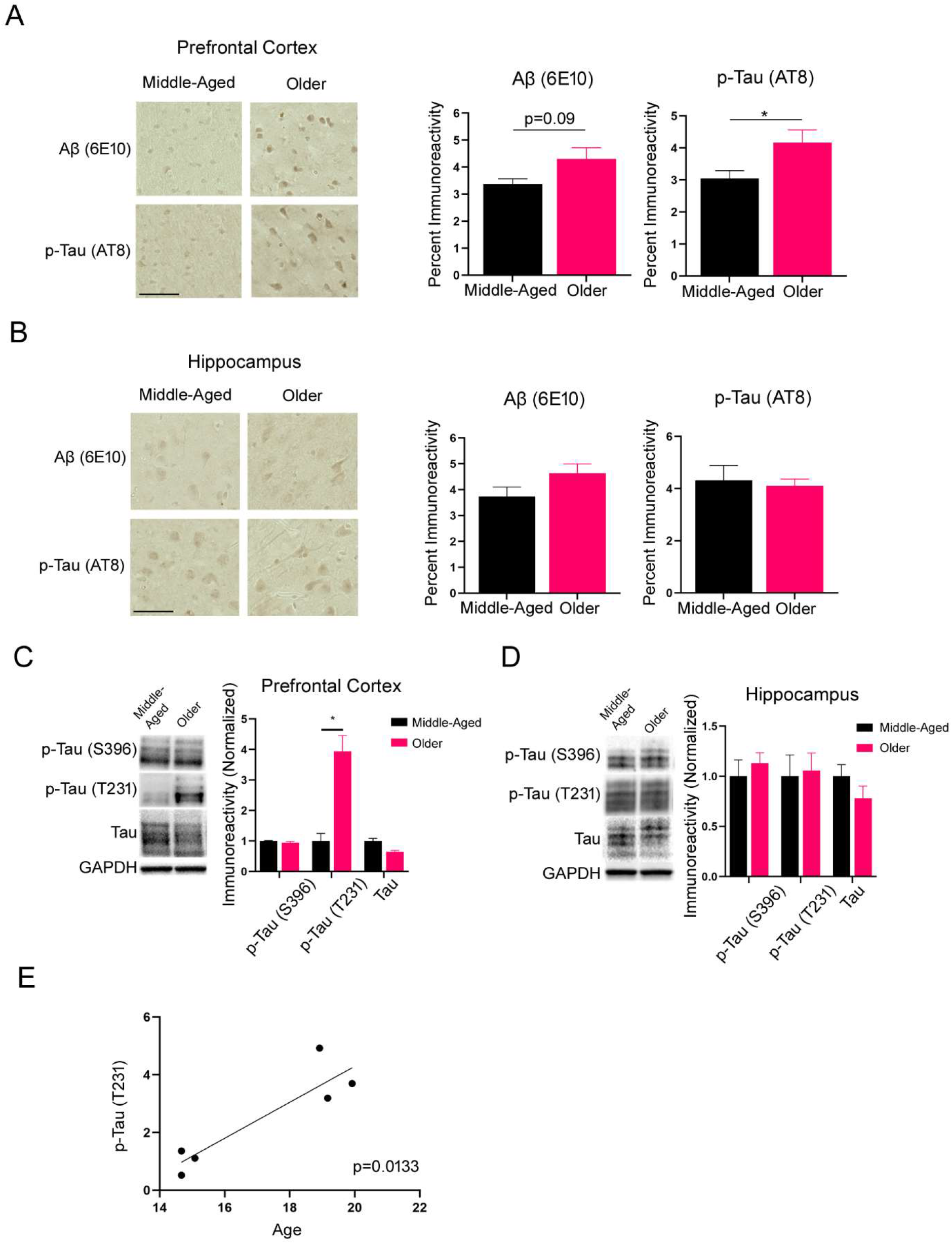
Increased tau phosphorylation in the prefrontal cortex of older NHPs. **(A)** Representative images and quantification of Aβ and p-tau immunostaining in the PFC of middle-aged and older monkeys. (Scale bar = 50 μm) **(B)** Representative images and quantification of Aβ and p-tau immunostaining in the hippocampus of middle-aged and older monkeys. (Scale bar = 50 μm). **(C)** Western blot analysis on tau phosphorylation in the prefrontal cortex of the NHPs. **(D)** Western blot analysis of tau phosphorylation in the hippocampus of the NHPs. Middle-Aged n = 3, Older n=3. \**p* < 0.05, unpaired *t* test. (E) Correlation between age and p-tau (Thr231) immunoreactivity in the prefrontal cortex. *p*< 0.05, Pearson correlation.

### 3.2. Age-related AMPK activation in the PFC of Cynomolgus macaques

Dysregulation of the AMPK signaling pathway has been associated with multiple AD pathophysiology based on studies from human samples and rodent models of AD (Domise et al., 2016; Vingtdeux et al., 2011; Wang, Xin et al., 2019; Zimmermann et al., 2020). Moreover, AMPK is considered as a tau kinase that can phosphorylate multiple sites in the microtubule-binding domain and within the flanking regions of tau protein including Ser396 and Thr231 (Augustinack et al., 2002; Domise et al., 2016; Thornton et al., 2011). By performing Western blot assay on the NHP brain tissue, we found that levels of phospho-AMPKα (Thr172), indicative of overall AMPK activity, were significantly increased in the PFC of older monkeys compared to the middle age group (Fig. 2A). Interestingly, levels of total AMPK protein in the PFC of older monkeys were reduced compared to middle age group (Fig. 2A). On the other hand, we did not observe age-related alterations on levels of AMPK phosphorylation or total AMPK protein in the hippocampus of older NHPs (Fig. 2B). Moreover, we examined alterations of glycogen synthase kinase 3 β (GSK3β), another established tau kinase that is implicated in AD pathogenesis (Hernandez et al., 2013; Sperber et al., 1995; Wagner et al., 1996). GSK3β activity, assayed by its phosphorylation at the Ser9 site, was analyzed with Western blot. No age-related change in levels of either GSK3β phosphorylation or total GSK3β protein was observed in PFC or hippocampus of the NHPs (Fig. 2C and D). Taken together with the above findings on brain tau pathology (Fig. 1), these data suggest a potential link between AMPK overactivation and tau hyperphosphorylation in PFC of NHPs.

**Figure 2.**
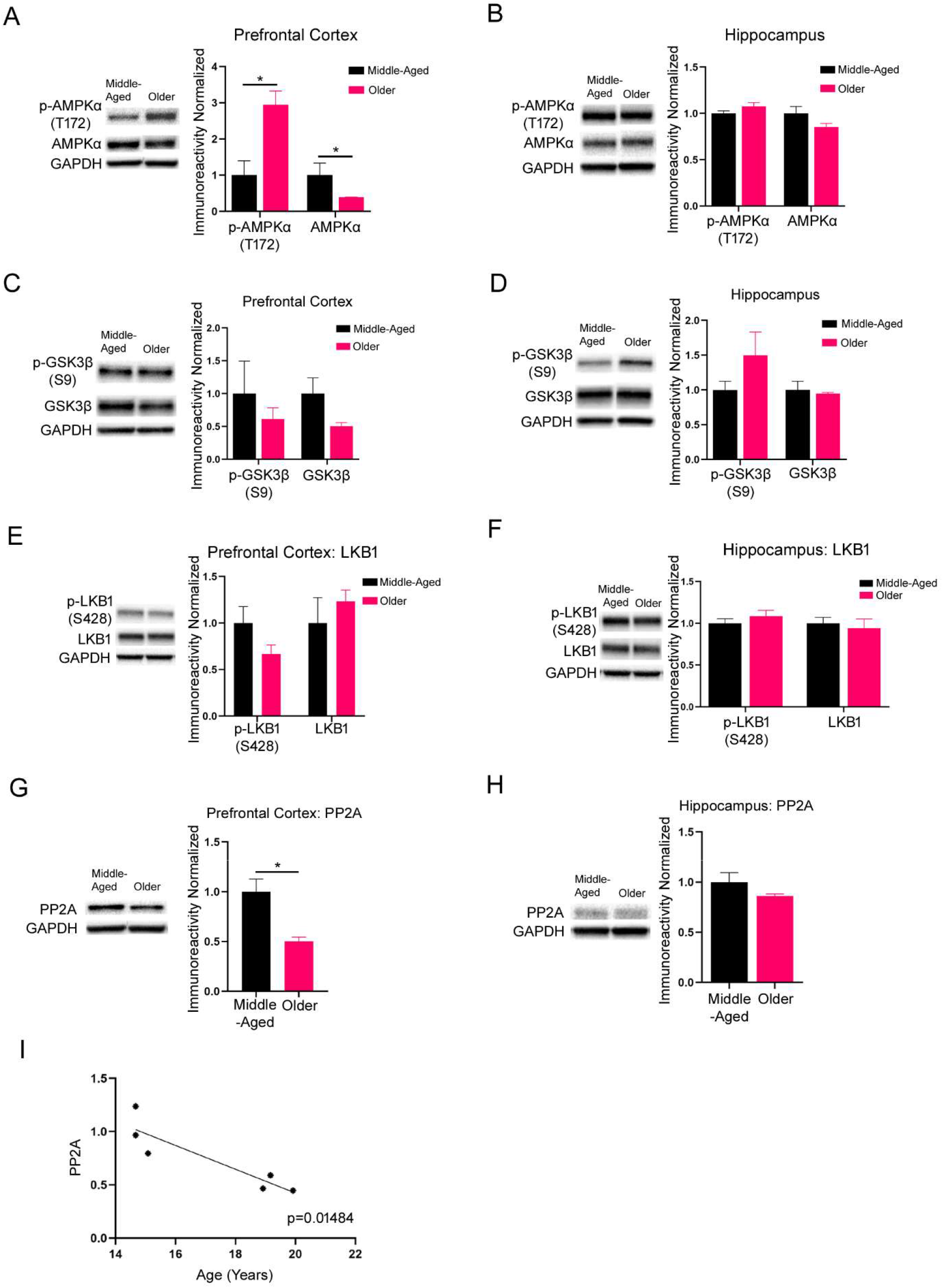
Increased AMPK activity in the prefrontal cortex of older NHPs. **(A)** Western blot analysis showed increased AMPK activity in the prefrontal cortex of older NHP group. **(B)** No change in AMPK activity in the hippocampus. **(C-D)** No change in GSK3β activity in the PFC or hippocampus of older monkeys. **(E-F)** No change in LKB1 activity in the PFC or hippocampus of older monkeys. **(G)** Decrease in PP2A protein levels in the prefrontal cortex of older group. Middle-Aged n = 3, Older n=3. \**p* < 0.05, unpaired t test. **(H)** No change in PP2A expression in the hippocampus. (**I**) Correlation between age and PP2A protein levels in the prefrontal cortex of the NHPs. *p* < 0.05, Pearson correlation.

To understand the molecular mechanisms underlying age-related AMPK phosphorylation, we analyzed the activity of an established AMPK kinase: liver kinase B 1 (LKB1) (Hardie et al., 2012; Yan et al., 2018). Surprisingly, we did not find any age-related alterations in activity of LKB1 in either PFC or hippocampus of the NHPs (Fig. 2E and F)

We next examined regulation of protein phosphatase 2A (PP2A), which functions as the main phosphatase for both AMPK and tau (Martin et al., 2013). Compared to the middle age NHPs, there was a significant decrease in PP2A protein levels in the PFC of the older group (Fig. 2G). No age-related alterations in PP2A levels were observed in the hippocampus of NHPs (Fig.2H). In addition, the decreased PP2A expression in the PFC correlated significantly with age (Fig. 2I). These findings indicate that abnormal de-phosphorylation activities (by PP2A) may contribute to the increased phosphorylation of tau and/or AMPK seen in the PFC of the older NHPs. In addition, we examined two downstream targets of AMPK: Acetyl-CoA carboxylase (ACC1) and Unc-51 like autophagy activating kinase (ULK1). Autophagy dysfunction has been implicated in AD and is negatively regulated by AMPK through inhibition of ULK1 (Egan et al., 2011; Hardie et al., 2012; Li et al., 2017). We observed no significant changes in AMPK-dependent phosphorylation of ACC1 or ULK1 (Supplemental Fig. 1A and B).

### 3.3. Age-related alterations of mitochondrial morphology and biology in the brain of NHPs

AMPK is a key regulator of mitochondrial biogenesis and dynamics through peroxisome proliferator-activated receptor γ coactivator 1-α (PGC-1α) signaling (Hardie et al., 2012). Previous studies with human samples and rodent models have shown defects of mitochondrial function in multiple neurodegenerative diseases including AD (Martín-Maestro et al., 2017; Wang, W. et al., 2020). Using transmission electron microscopy (TEM), we analyzed changes in mitochondrial morphology. We observed a significant increase in the overall number of mitochondria in the PFC of the older NHPs, compared to the middle age group (Fig. 3A). There was no change in average mitochondrial length and size in PFC of the older group compared to middle-aged monkeys (Fig. 3A). In comparison, we found significant decreases in the average length and size of mitochondria in the hippocampus of the older group without alterations in overall hippocampal mitochondrial number (Fig. 3B). We further analyzed expression of key proteins in mitochondrial biogenesis and dynamics including PGC-1α, mitofusin2, and dynamin related protein 1 (DRP1). None of these proteins showed age-related alterations in either the PFC or hippocampus of the NHPs, except for a trending increase of PGC-1α protein levels in the PFC of the older group (Fig. 3C and D). To further determine potential age-related changes in mitochondrial function, we analyzed regulation of the membrane-bound protein complexes that compose the ETC of mitochondria. These complexes are responsible for the majority of ATP production and are dysregulated in AD (Maruszak and Żekanowski, 2011; Reddy and Beal, 2008). The OXPHOS antibody cocktail recognizes a key subunit from each of the five complexes in the ETC. We found no aging-related changes in the PFC and only a trending (p=0.1) decrease in complex IV subunit expression in hippocampus of older NHPs (Supplemental Fig. 2A and B). Dysfunctional mitochondria is associated with oxidative stress, and we further analyzed potential age-related alterations of superoxide dismutase (SOD) 1 and 2, and catalase expression. SOD1 and 2 function to clear reactive oxygen species (ROS) (Cao et al., 2018; Wojsiat et al., 2018). Catalase is an enzyme that breaks down the hydrogen peroxide created by SOD1 and 2 and is also integral in clearing harmful ROS (Wojsiat et al., 2018). There was a significant decrease in SOD1 expression in the PFC of the older group with no other changes of SOD1/2 or catalase observed in PFC or hippocampus (Supplemental Fig. 3A and B). Taken together, these findings indicate mitochondrial deficiency in NHPs during transition from middle age to old age.

**Figure 3.**
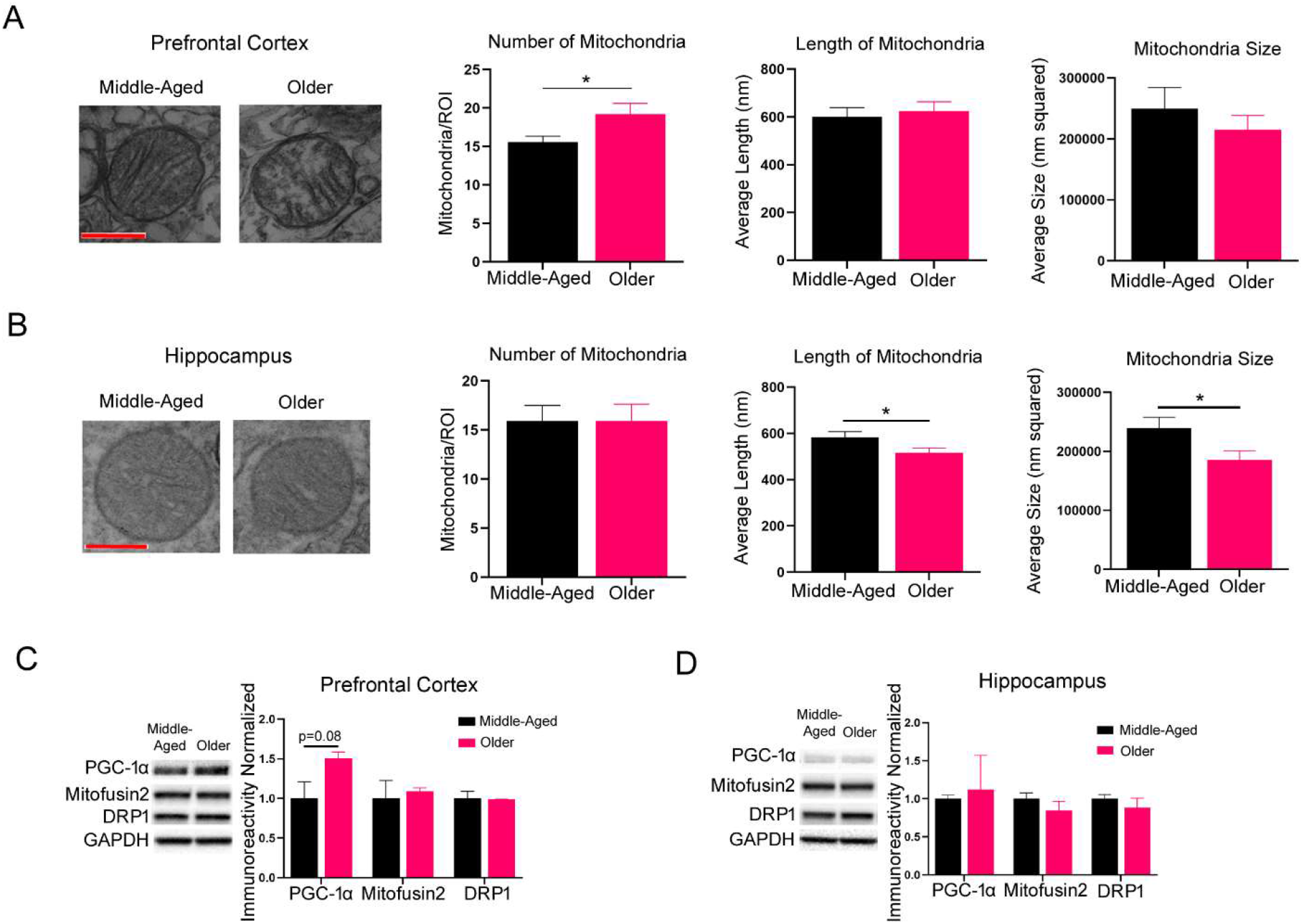
Impaired mitochondrial morphology in PFC and hippocampus of NHPs with age. **(A)** Analysis of mitochondrial morphology in PFC of the NHPs. Representative TEM images of mitochondria in the PFC were shown. (Scale bar = 500nm). Significant increase in the number of mitochondria in the older group, and no change in mitochondria length or size in the PFC. Middle-Aged n = 3, Older n=3; 20 ROIs per animal. \**p* < 0.05, unpaired t test. **(B)** Analysis of mitochondrial morphology in hippocampus of the NHPs. Representative TEM images of mitochondria in the hippocampus were shown. (Scale bar = 500nm). No change in the number of mitochondria in the older group in the hippocampus. Significant decrease in mitochondria length and size in the hippocampus of older group. Middle-Aged n = 3, Older n=3; 20 ROIs per animal. \***p** < 0.05, unpaired t test. **(C)** Trending increase in PGC-1α expression, but no change in expression of Mitofusin2 or DRP1 in the PFC of older group via Western blot quantification. **(D)** No change in PGC-1α, Mitofusin2 or DRP1 expression in the hippocampus of older group.

### 3.4. Age-related alterations in postsynaptic density of NHPs

AD is considered as a disease of “synaptic failure” (Selkoe, 2002). Dendritic spine loss and impairments of synaptic plasticity have been implicated in the early stage of AD (Forner et al., 2017; Knobloch and Mansuy, 2008; Overk and Masliah, 2014; Scheff et al., 2006). The postsynaptic density (PSD) is an integral part of the synapse and plays a critical role during synaptic transmission (Sheng and Kim, 2011). Using TEM, we analyzed potential age-related alterations of PSD morphology in the NHPs. In the PFC, we observed a significant decrease in both the number and size of PSDs in the older NHPs, compared to the middle age group (Fig. 4A). In the hippocampus there were no difference in number of PSDs or in the length of the active zone between the two groups (Fig. 4B). Interestingly, we found a nearly significant increase in the average size of PSDs in the hippocampus of the older NHPs, compared to the middle age group (Fig. 4B). Additionally, there were no significant changes in PSD-95 or Synapsin2 protein expression levels in either the PFC or hippocampus of the older group (Supplemental Fig. 4A and B). In addition, synaptic alterations are often reliant on protein synthesis mechanisms. Mammalian target of rapamycin (mTOR) is a key regulator of protein synthesis with several proteins in the signaling cascade being downstream of AMPK (Wang, X. et al., 2019). We examined the activities and expression levels of several mTOR signaling molecules including mTOR1, S6K1, 4E-BP1, and eEF2 but found no significant changes in either the PFC or hippocampus of the older group compared to the middle-aged group (Supplemental Fig. 5A and B).

**Figure 4.**
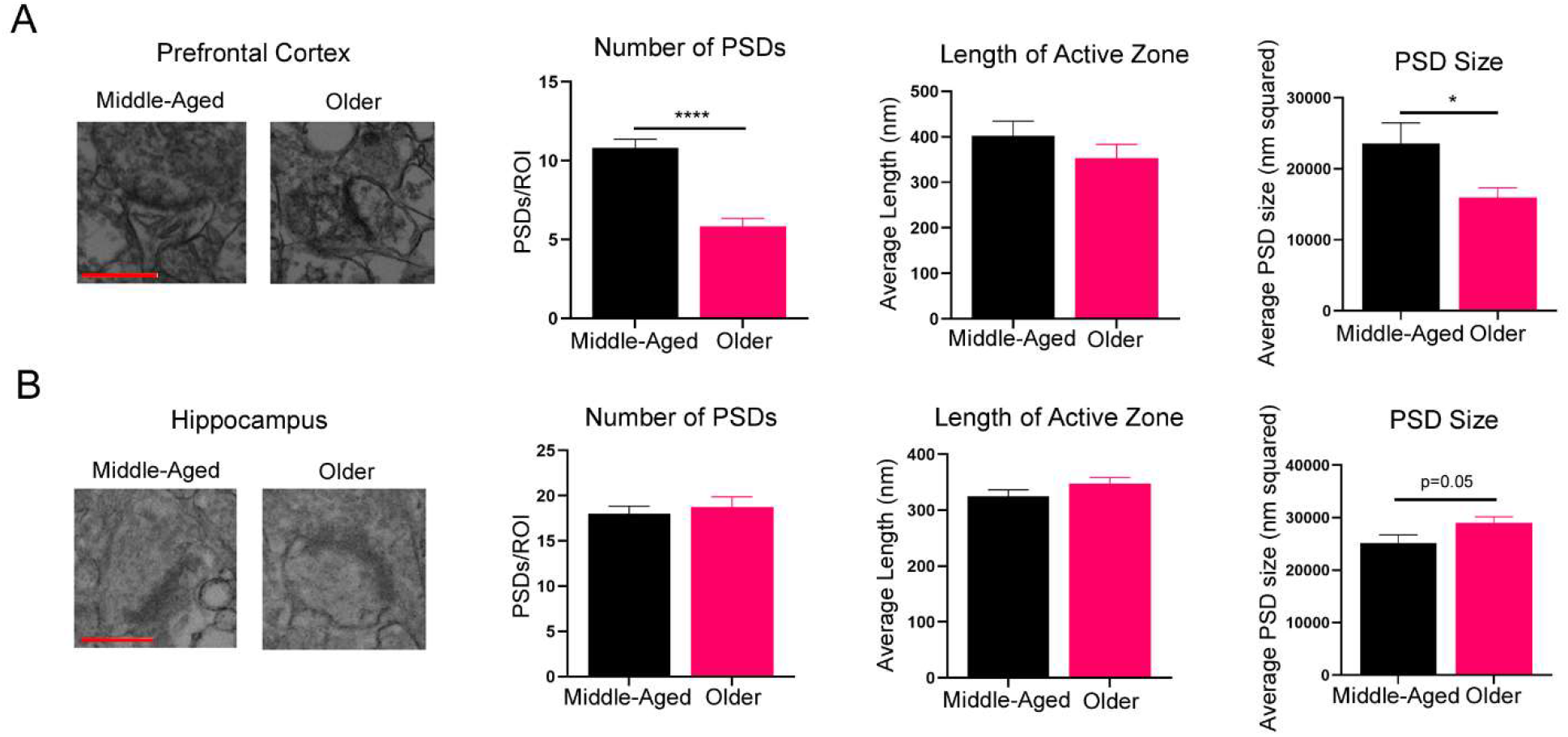
Age-related alterations of postsynaptic density in the brain of NHPs. **(A)** Analysis of PSDs in the PFC of NHPs. Representative TEM images of PSDs in the PFC were shown. (Scale bar = 500nm). Significant decrease in the number of PSDs in the PFC of the older group, and no change in length of active zone in the PFC with age. Decrease in average PSD size in the PFC of monkeys with age. Middle-Aged n = 3, Older n=3. 20 ROIs per animal. \**p* < 0.05, \*\*\*\**p* < 0.0001, unpaired *t* test. **(B)** Analysis of PSDs in the hippocampus of NHPs. Representative TEM images of PSDs in the hippocampus were shown. (Scale bar = 500nm). No change in the number of PSDs or the length of the active zone in the hippocampus of the older. Increase in average PSD size in the hippocampus of older group. Middle-Aged n = 3, Older n=3. 20 ROIs per animal. Unpaired t test.

## 4. Discussion

Majority of the current rodent AD models are generated via genetic manipulation mimicking FAD, which is characterized by definite genetic cause and early onset of cognitive syndromes (Puzzo et al., 2015). Meanwhile, sporadic AD or late onset AD (LOAD) accounts for more than 90% of all AD cases, with aging the most known risk factor. Recent studies carried out in different species of aged NHPs demonstrated pathological features in the animals resembling those in human LOAD (Arnsten et al., 2019; Cramer et al., 2018; Latimer et al., 2019; Oikawa et al., 2010; Paspalas et al., 2018; Van Dam and De Deyn, 2017). Given their close phyletic relationships with human beings, findings from the study of NHPs could provide important insights into AD pathogenesis that might be unrecognized with research in FAD models. (Arnsten et al., 2019). In this pilot study, we focused on examining potential AD-like pathological alterations in Cynomolgus macaques during their transitioning from middle age (14-15 years old, equivalent to 48 year old in human) to “older” age (18-19 years old, equivalent to 62 year old in human). Previous studies in old Cynomolgus macaques (>20 year old) have found evidence of Aβ plaques in frontal and temporal cortices (Nakamura et al., 1998; Wu et al., 2008). In comparison, we observed hyperphosphorylated tau (assessed by AT8 staining) pathology but no Aβ plaques in the brain of older monkeys (Fig. 1). This raises an interesting question whether tau pathology occurs before Aβ plaque accumulation in the brain with aging and/or during development of AD. While substantial studies have demonstrated that Aβ accumulation results in tau hyperphosphorylation (Small and Duff, 2008), accumulating evidence suggests a more complicated relationship between Aβ and tau pathology during development of AD, particularly LOAD (Arnstern, 2020). Neuropathological analysis of brain samples from both human and NHPs indicates that abnormal tau phosphorylation may happen prior to Aβ accumulation in the brain, triggering multiple pathological events that lead to synaptic failure and cognitive deficits later in life (Braak and Del Tredici, 2011; Braak et al., 2011; Darusman et al., 2014; Oikawa et al., 2010). It is worth mentioning that most of the AD-like pathological features in older monkeys displayed in PFC but not hippocampus, which is different from studies done in rodent AD models (Elder et al., 2010; Heilbroner and Kemper, 1990; Mufson et al., 1994; Poduri et al., 1994; Sani et al., 2003). Research on vulnerability of different brain regions and/or cell types to AD has drawn much attention recently, and future more comprehensive studies in NHPs are warranted to elucidate the basis for such regional (and probably cellular) vulnerability.

Previous studies in rodents showed that over-activation of AMPK leads to impairments of long-term potentiation (LTP), a cellular model for memory and major form of synaptic plasticity (Potter et al., 2010). Further, elevated AMPK activity has been associated with multiple aspects of AD pathophysiology. Consistently, inhibiting the activities of AMPK and its downstream effectors alleviates synaptic failure and cognitive deficits in transgenic mouse models of AD (Domise et al., 2016; Wang, Xin et al., 2019; Zimmermann et al., 2020). In agreement with these reports, we observed increased AMPK activity (measured by p-AMPKα) in the PFC (but not hippocampus) of older monkeys (Fig. 2A). Interestingly, our data showed a marked correlation between increased phosphorylation of AMPK and decreased protein expression of PP2A, a phosphatase with a broad spectrum of substrates including AMPK and tau (Martin et al., 2013; Salminen et al., 2016). While much efforts have been dedicated to therapeutic strategies targeting kinases, roles of phosphatases (particularly PP2A) in AD etiology have drawn a lot of attention recently (Leong et al., 2020; Nicholls et al., 2016; Torrent and Ferrer, 2012; Wei et al., 2020).

There are limitations for the current study. This is a pilot study with limited number of subjects (n=3 monkeys for each group). The monkeys in the older group also exhibit slightly higher (not significant statistically) levels of blood glucose and triglycerides (Table 1), which may contribute to the AD-like pathologies described in the results. Additionally, all three monkeys in the older group are female, while the middle age group includes two female and one male monkeys. We hope to accumulate more aged macaque subjects (with middle age controls) in the future to dissect out potential “confounding” factors for the current study. Further, diagnosis of AD in clinical practice requires evidence showing pathological and biochemical changes in the context of dementia syndromes. Unfortunately, no data on behavioral assessment are available for the monkeys in this study. Future in-depth research in NHPs, particularly longitudinal studies with functional assessment, is necessary to characterize how the aforementioned pathological alterations are involved in the development of neurodegeneration and cognitive syndromes in AD, thus providing insights into potential diagnostic/prognostic biomarkers and therapeutic/preventive strategies for this devastating human disease.

## Disclosure statement

The authors declare no conflict of interest.

## Acknowledgements

We thank the Wake Forest School of Medicine Pathology and Imaging Core for their help with tissue processing and technical help with imaging experiments. This work was supported by National Institutes of Health grants R01 AG055581, R01 AG056622 (T.M.), the Alzheimer’s Association grant NIRG-15-362799 (T.M.), the BrightFocus Foundation grant A2017457S (T.M.).

---

**Supplemental Figure 1.**
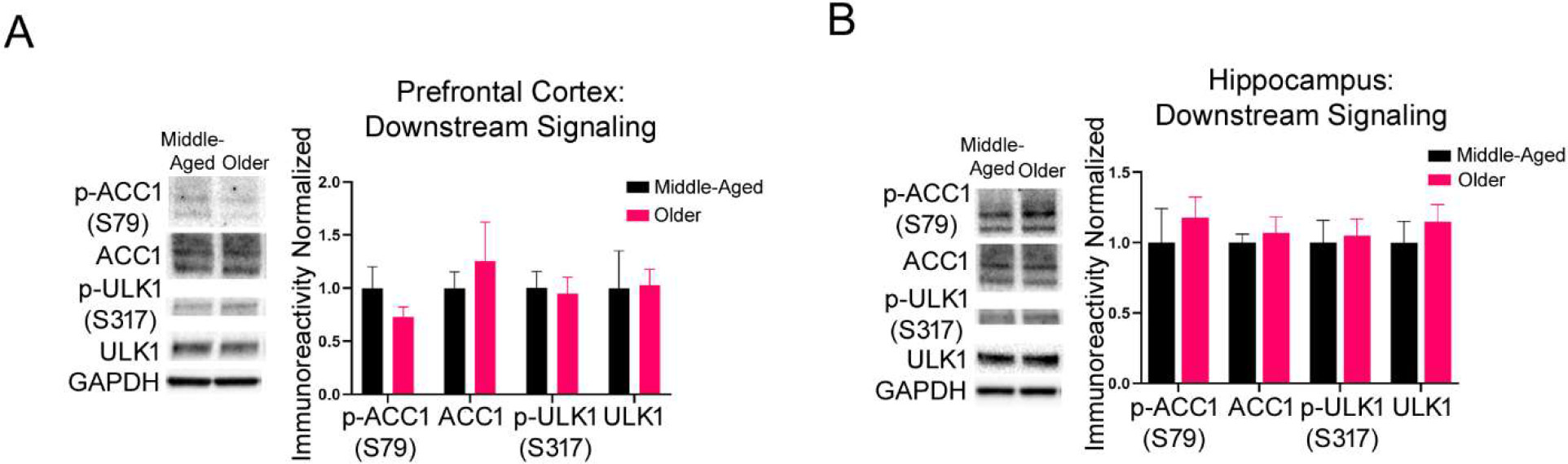
Examination of molecules downstream of AMPK signaling. **(A)** Western blot analysis on activities of ACC1 or ULK1 in the PFC of NHPs. **(B)** Western blot analysis on activities of ACC1 or ULK1 in the hippocampus of NHPs. Middle-Aged n = 3, Older n=3.

**Supplemental Figure 2.**
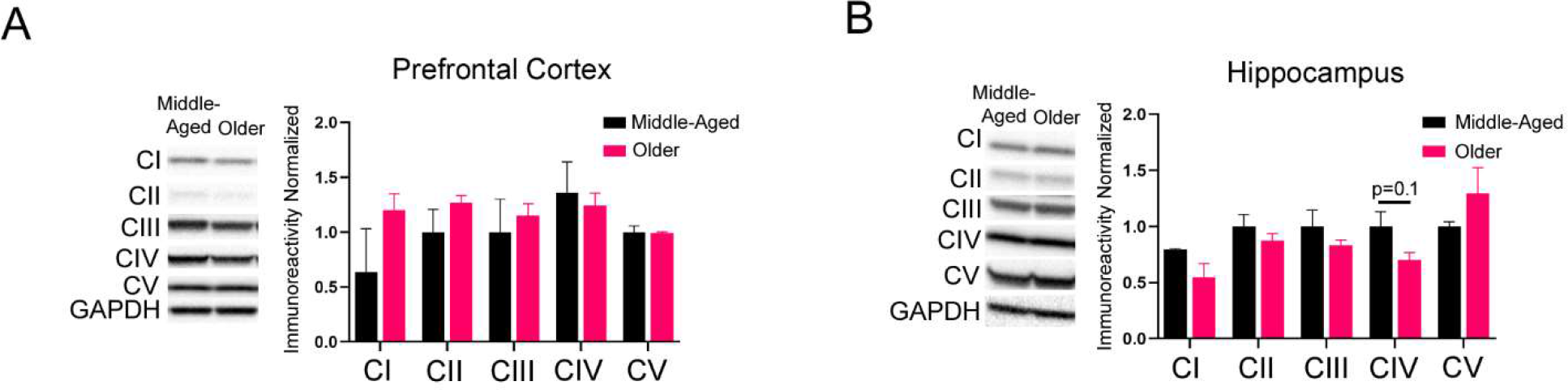
OXPHOS complex expression in PFC (A) and hippocampus (B) of NHPs. Middle-Aged n = 3, Older n=3.

**Supplemental Figure 3.**
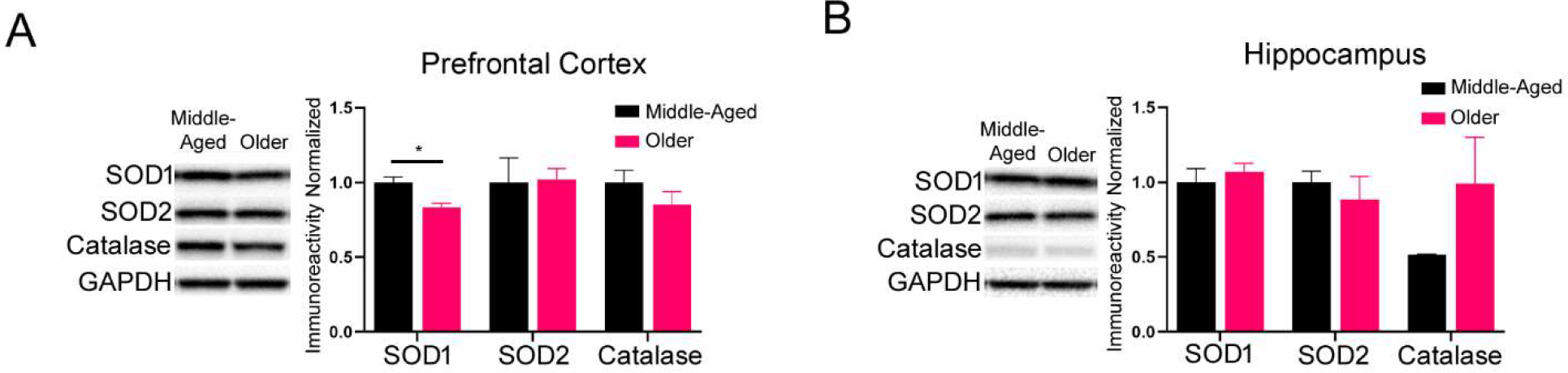
Analysis of proteins involved in clearance of reactive oxygen species (ROS) in NHPs. (**(A)** Western blot assays on protein levels of SOD1/2 and Catalase in PFC. (**B**) Western blot assays on protein levels of SOD1/2 and Catalase in hippocampus. Middle-Aged n = 3, Older n=3. \**p* < 0.05, unpaired t test.

**Supplemental Figure 4.**
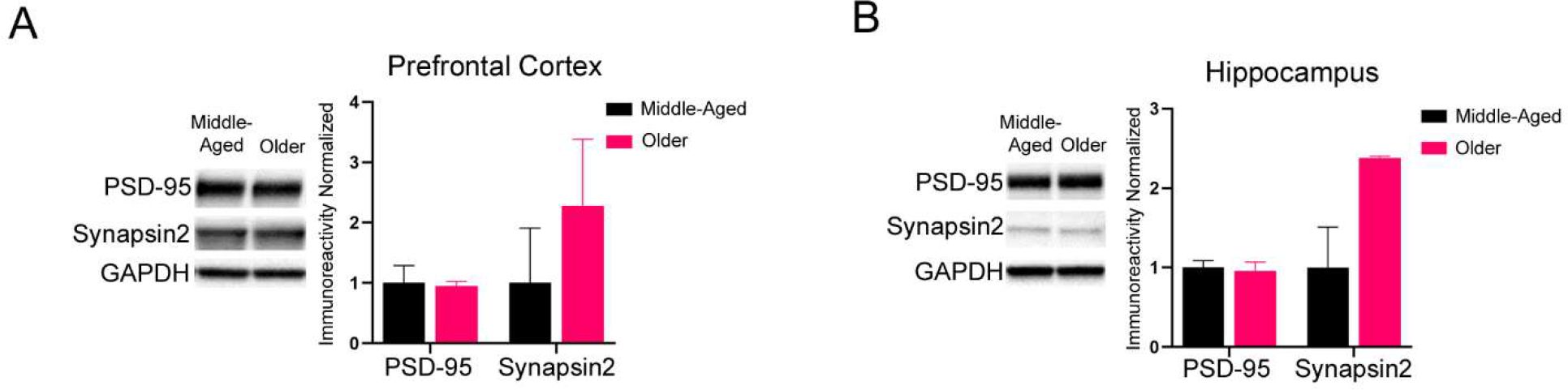
Examination of synaptic proteins in NHPs. **(A)** Western blot analysis of the synaptic proteins PSD-95 and Synapsin2 in PFC. (**B**) Western blot analysis of the synaptic proteins PSD-95 and Synapsin2 in hippocampus. Middle-Aged n = 3, Older n=3.

**Supplemental Figure 5.**
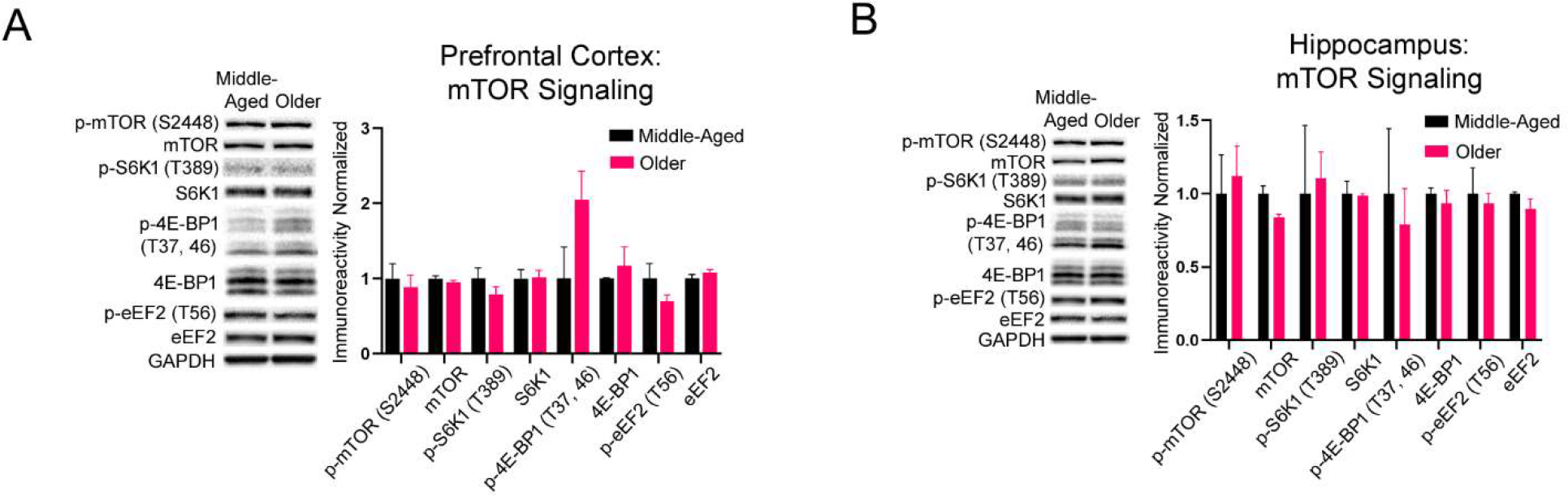
Analysis of mTOR signaling in NHPs. (**A**) Western blot analysis of on activities of mTOR, S6K1, 4E-BP1, and eEF2 proteins in PFC. (**B**) Western blot analysis of on activities of mTOR, S6K1, 4E-BP1, and eEF2 proteins in hippocampus. Middle-Aged n = 3, Older n=3.

## Notes

### Competing Interest Statement

The authors have declared no competing interest.

